# Investigations on regulation of miRNAs in rice reveal [Ca^2+^]_cyt_ signal transduction regulated miRNAs

**DOI:** 10.1101/2021.03.24.436297

**Authors:** Shivani Kansal, Vaishali Panwar, Roseeta Devi Mutum, Saurabh Raghuvanshi

**Author notes:** **Corresponding author**: Dr. Saurabh Raghuvanshi, Department of Plant Molecular Biology, University of Delhi South Campus, Benito Juarez Road, New Delhi – 110021, India. **Authors’ email addresses**.

## Abstract

MicroRNAs are critical components of the multi-dimensional regulatory networks in eukaryotic systems. They regulate a spectrum of developmental and metabolic processes in both plants and animals. Thus, it is quite apparent that the transcription, processing as well as activity of the miRNAs themselves is very dynamically regulated. One of the most important and universally implicated signalling molecule is [Ca^2+^]_cyt_. It is known to regulate a plethora of developmental and metabolic processes in both plants and animals, however their impact on the regulation of miRNA expression is relatively less explored. The current study employed a combination of internal and external calcium channel inhibitors, to establish that [Ca^2+^]_cyt_ signatures actively regulate miRNA expression in rice. Involvement of [Ca^2+^]_cyt_ in regulation of miRNA expression was further confirmed by treatment with calcimycin, the calcium ionophore. Modulation of the cytosolic calcium levels was also found to regulate the drought responsive expression as well as ABA mediated response of miRNA genes in rice seedlings. The study further establishes the role of calmodulins and Calmodulin-binding Transcription Activators (CAMTAs) as important components of the signal transduction schema that regulates miRNA expression. Yeast-one-hybrid assay established that OsCAMTA4 & 6 are involved in the transcriptional regulation of miR156a and miR167h. Thus, the study was able to clearly establish that [Ca^2+^]_cyt_ is actively involved in regulating expression of miRNA genes both under control and stress conditions.

## Introduction

Plants being sessile in nature need to face all the environmental and biotic challenges at one place. Plants modulate their internal system to cope with these challenges. Changes in the growth or environment of the plants is conveyed to the cell via primary messengers such as hormones while it is relayed to the nucleus via the secondary messengers such as cAMP, cGMP, Inositol triphosphate, diacylglycerol and the most widely studied [Ca^2+^]_cyt_. Apart from these, players like hydrogen sulphide, hydrogen peroxide, carbon monoxide and nitric oxide collectively termed as ‘gasotransmitters’ have also emerged as potential signalling messengers in cell (Lisjak et al., 2013; Mittler et al., 2011; Gaupels et al., 2011). Their signalling cascade is often routed via [Ca^2+^]_cyt_ or MAPK signal transduction components. [Ca^2+^]_cyt_ as a second messenger of internal and external signals has been well studied and documented in a plethora of studies and reviews describing its role in signalling in various plant physiological functions such as abiotic stress response (Dodd et al., 2010), stomatal aperture (McAinsh et al., 1990), self-incompatibility (Franklin-Tong et al., 1993), pathogenic and symbiotic interactions (Ma & Berkowitz, 2007) and growth of pollen tube and root tips (Hepler et al., 2001). Upon perceiving an environmental stimulus or developmental cue, rapid and precise [Ca^2+^] oscillations are generated within the cytoplasm through influx of [Ca^2+^] from internal as well as extracellular stores. Besides cytosolic [Ca^2+^] signatures and signalling, the plant cell nucleus has also been proved to contain within an independent Ca^2+^signalling system (Pauly et al., 2001; Pauly et al., 2000; Tou et al., 2004) to regulate gene transcription. Calmodulin (CaM) is said to be the primary decoder of the signal along with the support of CaM-binding proteins in the nucleus. For instance, nuclear apyrase in pea binds to CaM and is activated by Ca^2+^/CaM complex (Hsieh et al., 2000); nuclear localized potato CaM-binding protein (PCBP) identified from developing potato tubers binds to CaM in Ca^2+^ dependent manner (Reddy et al., 2002). The strongest evidence for the theory is the identification of nuclear transcription factors that are regulated directly or indirectly by CaM (Yang & Poovaiah, 2003). For instance, CAM7 in *Arabidopsis* has been demonstrated to regulate photomorphogenic growth and light-responsive gene expression by binding to Z-/G-box elements in their promoters including CAB1 and RBCS1A. CAM7 was also shown to work independently yet in an inter-dependent pathway with HY5 to regulate the photomorphogenic growth of *Arabidopsis* (Kushwaha et al., 2008). WRKY class of TFs has also been shown to interact with Ca^2+^/CaM complex especially the WRKY group IId (Park et al., 2005). Another separate class of transcription factors known as CAMTAs (Calmodulin-binding transcription activators), described to be evolutionary conserved from humans to plants (Finkler et al., 2006) respond rapidly to multiple abiotic stresses such as drought, salinity, heat, cold, UV thus regulating the signalling required (Yang & Poovaiah, 2003). Despite the name, CAMTAs can act as transcriptional activators as well as repressors; as CAMTA3 was demonstrated for activating the expression of *CBF2* imparting cold tolerance (Doherty et al., 2009) as well as repress the expression of *EDS1*, a regulator of salicylic acid level (Du et al., 2009). Recently, AtCAMTA3 was shown to repress the genes involved in SA-biosynthesis and SA-mediated immunity in healthy plants (Kim et al., 2017).

In biological systems miRNAs represent a significant and critical layer of regulation of protein coding genes at the post-transcriptional level. In our previous studies and others as well, we have seen that several miRNAs can express differentially to similar cue in different rice cultivars, thus emphasizing the evolution of distinct regulatory mechanisms controlling miRNA expression in different cultivars (S. Balyan et al., 2017; Kansal et al., 2015; Mutum et al., 2013). Such unique regulation would have a cascading effect on plant biology since a single miRNA regulates several protein coding genes. Thus, study of regulation of miRNA expression is of critical importance. MiRNAs have been shown to be differentially regulated by different stages of growth and development, biotic and abiotic stresses etc. At the molecular level, miRNAs have been shown to respond to secondary messengers of signalling in various publications. A study showed seven miRNAs including miR169, miR397, miR1425, miR408-5p and miR827 that were upregulated while miR319a-3p and miR408-5p that were down-regulated by H_2_O_2_ stress (T. Li et al., 2011). Similarly, miRNAs were also shown responsive to exogenously supplied H_2_S and it was established that their expression under drought simulation was mediated via H_2_S (Jiejie Shen et al., 2013). Analysis in the *lcd* (knockdown mutant of H_2_S producing enzyme L-CYSTEINE DESULFHYDRASE) mutants under control and drought simulated conditions showed lower expression levels of some miRNAs like miR167a/c/d, miR393a, miR396a, miR398a/b/c which could be rescued by resupplying H_2_S. This confirmed that H_2_S regulates miRNA activity to modulate the drought response of *Arabidopsis*. In contrast, to the best of our knowledge the demonstration that miRNAs are responsive to [Ca^2+^]_cyt_ has not been made till now.

In the current study, we attempt to establish a stronger link between calcium signalling and its control over regulation of miRNAs in rice. The miRNAs were confirmed to be calcium responsive using miRNA expression tools (small RNA libraries and qRT-PCR) under conditions of calcium scarcity (created using Ca^2+^ channel inhibitors) and excess of [Ca^2+^]_cyt_ using a Ca^2+^ ionophore A23187. Further, role of Ca^2+^ signalling during differential regulation of miRNAs under abiotic stress such as dehydration and hormonal stress such as ABA was also established. Additionally, we show here the involvement of calmodulins and CAMTAs in the regulation of miRNA expression. Furthermore, our results show physical interaction between OsCAMTA4 and OsCAMTA6 and the promoters of miR156a and miR167h thereby strengthening the evidence of hold of calcium signalling over miRNA regulation.

## METHODS

### Rice Plant Growth and Stress Treatment

Rice seeds [*O*.*sativa* subsp. *indica* var Nagina22 (N22)] were obtained from Indian Agricultural Research Institute (IARI) Pusa, New Delhi. Seeds were sterilized and planted as per Kansal *et al*. 2015, grown for 1 week and then subjected to respective stresses. For seedling stress experiments, the seedlings were transferred to a solution of calcium channel inhibitors [mix of LaCl_3_ (5mM), Verapamil (50μM), LiCl (5mM), Ruthenium red (100μM) for 3h], calcium ionophore A23187 (10μM for 5h), TriFluoPerazine (200 μM for 6h) or ABA (100μM for 3h). All these solutions and control (MQ water) contained the surfactant Silwett (Sigma) at 0.01% final concentration for better and uniform absorption. All the chemicals were obtained from Sigma Aldrich. For dehydration stress, the seedlings were taken out of the respective solution, dabbed with tissue paper to remove excess liquid sticking to the roots, and kept open in the air for air-drying until leaf rolling was observed. Drought stress at mature stage was given as per Mutum et al., 2016.

### RNA Isolation and cDNA synthesis

RNA was isolated using TriReagent (Sigma) as per the manufacturer’s protocol. It was subjected to DnaseI treatment followed by small RNA enrichment by 4 M LiCl method as given in (Kansal *et al*. 2015). For cDNA synthesis, 2 μg of RNA was incubated with 100 pmoles of either oligodT primer (Eurofins Genomics) for total RNA or miR_oligodT primer (Eurofins Genomics) for small RNA using the SuperscriptII cDNA synthesis kit (Invitrogen). All the primers have been enlisted in Supplementary Table S1.

### Small RNA Library Preparation, quality check and analysis

Small RNA libraries were prepared with 2 μg of enriched small RNA using NEBNext® Small RNA Library Prep Set for Illumina® (Multiplex Compatible) (New England Biolabs) according to the manufacturer’s protocol in biological dupicates. The libraries thus prepared were checked for quality using 2100 Bioanalyzer (Agilent) High Sensitivity DNA chip according to the manufacturer’s protocol. The libraries were sequenced using Illumina platform by Sandor LifeSciences Pvt Ltd. Small RNA Library Analysis was done using CLC Genomics Workbench version 7.0.4 (Qiagen) with the help of ‘Small RNA Analysis’ tool. For further detailed analysis of differentially expressed genes the ‘Statistical Analysis’ tool, in particular ‘Empirical Analysis of DGE’ that uses ‘Edge’ as the tool for calculating differentially expressed genes, was used.

### Quantitative Real Time PCR

The first strand cDNA synthesized was used for qRT-PCR with appropriate dilution. The reaction was set up by adding Taqman Master Mix chemistry (2x). The reaction was run in ABI Step-One Plus Real Time system as per its default cycling for Taqman chemistry. Rice actin and 5S rRNA was used as endogenous controls for a protein coding gene and miRNA profiling, respectively. For each sample, at least three technical and three biological replicates were set up. The use of * above a bar in the graph depicts its significance as * p≤0.05 as recorded by t-test.

### miRNA-Target analysis and classification

Putative targets of miRNAs were selected from in-house analysed publicly available rice degradome libraries (GSM455939, GSM455938, GSM434596, GSM960648, GSM476257; (Li et al., 2010; Wang et al., 2012; Wu et al., 2009; Zhou et al., 2010) as well as in-house generated three additional degradome libraries from spikelets at heading and anthesis stage, flag leaf at heading stage of N22 (Mutum et al., 2016). The targets falling within the criteria of category 0-2 along with read no ≥ 10 and pval≤0.05 were selected as putative targets. The target loci were then used to extract their GO annotations from RGAP database and classified according to their GO term of ‘molecular function’. Similarly, the loci were checked for their presence in the Ricecyc database available at Gramene database.

### Yeast one hybrid assay

Yeast one hybrid assay was performed using EZ yeast transformation kit (MP Biomedicals) as per the instructions. 110 bp, 80 bp and 200 bp region of miR156a promoter, miR168a promoter and miR167h promoter, respectively, was PCR amplified from the genomic DNA of N22 and cloned into pAbAi vector as bait, harboring the aureobasidin (AUR1-C) reporter gene. Full-length CDS of *OsCAMTA4* and *OsCAMTA6* was PCR amplified from genomic DNA of N22 and cloned into the pGADT7-AD vector as prey. The miR156a-pAbAi, miR167h-pAbAi, OsCAMTA4-pGADT7 and OsCAMTA6-pGADT7 plasmids were linearized and co-transformed into the Y1H gold yeast strain. The transformants were selected on SD medium lacking Uracil (U) and Leucine (L) (SD/-L-U), respectively, as per the manufacturer protocol. Colonies obtained were screened for positive insert via colony PCR. Further, the positive clones were inoculated as primary culture and subsequently used in the secondary culture (3 mL) and grown at 30°C, 200 rpm for 3 h till the OD = 1 at 600 nm. Each culture was then serially diluted (10^−1^, 10^−2^ and 10^−3^) and droplets of 10μL of each dilution including the undiluted culture were placed on the selection media (SD/-L-U) and incubated at 30°C for 3–5 days till the formation of colonies. The interaction was determined based on the ability of co-transformed yeast cells to grow on dropout media lacking Uracil and Leucine with 100, 200 and 300 ng/ml of Aureobasidin A.

## Results

### 1) miRNAs are responsive to [Ca^2+^]_cyt_ in N22 rice

Cellular calcium levels are primarily regulated by the activity of the calcium channels present either on the plasma membrane (blocked by LaCl_3_ and Verapamil) and/or the mitochondrial (inhibited by Ruthenium red) or endoplasmic reticulum membrane (blocked by LiCl). Thus, tweaking the activity of these membrane channels with the help of calcium channel inhibitors would lead to a change in the cellular calcium levels. In order to identify miRNA genes that are regulated by [Ca^2+^]_cyt_ a cocktail of Ca^2+^ channel blockers (viz. LaCl_3_, LiCl, Ruthenium Red and Verapamil) was used to treat young rice seedlings followed by sequencing of the sRNA population. The efficiency of the inhibitor treatment was confirmed by monitoring the expression of a calcium responsive ‘Ca^2+^ dependent protein kinase’ OsCPK6 (Wan et al., 2007) (Fig. S1). Briefly describing, 293 and 303 miRNAs were detected in control libraries while 258 and 279 miRNAs were detected from the tissue treated with calcium channel inhibitors (Supplementary Table S2). Further comparative analysis of the datasets revealed 17 differentially expressed miRNAs (fold change ≥ 2 and *p*≤0.05) (Supplementary Table S3), out of which 10 were found to be down-regulated while 7 were up-regulated in the presence of Ca^2+^ channel inhibitors (Fig. 1a). Subsequently, qRT-PCR analysis confirmed the calcium dependent expression of 11 miRNAs in rice (miR1425-5p, miR156a, miR159b, miR166g-3p, miR167h-5p, miR1862a,d, miR1876, miR1878, miR396c-3p, miR444b.1) (Fig. 1b).

**Fig. 1.**
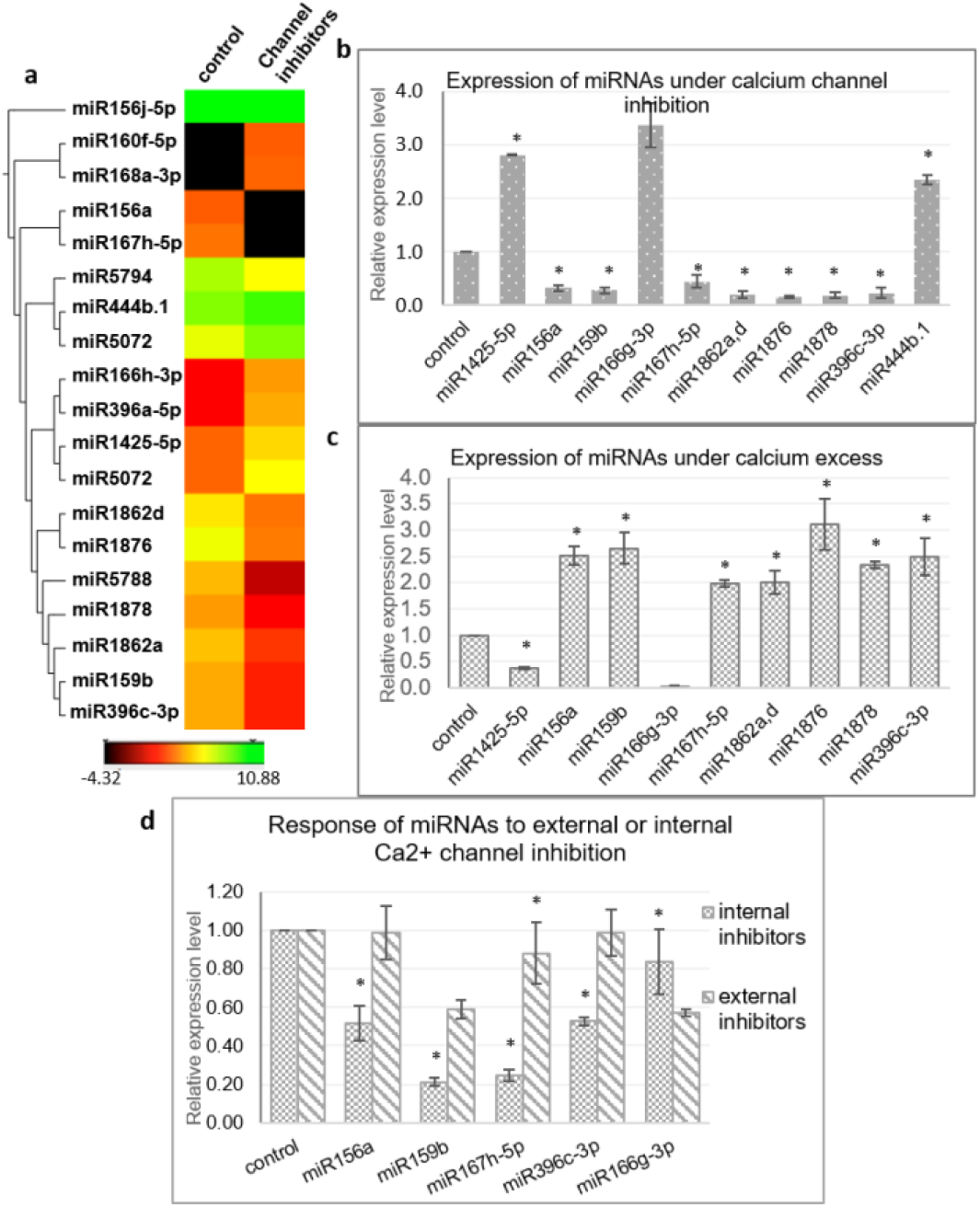
Identification of [Ca^2+^]_cyt_ responsive miRNAs in N22. **a** Hierarchical clustering of miRNAs detected with significant fold change in presence of [Ca^2+^]_cyt_ channel inhibitors. **b**. Validation of [Ca^2+^]_cyt_ responsiveness of selected miRNAs by qRT-PCR analysis. **c** Confirmation of [Ca^2+^]_cyt_ responsiveness by qRT-PCR generated expression profiles under calcium ionophore treatment. **d** qRT-PCR generated expression profiles of select miRNAs under treatment with specific set of Ca^2+^ channel blockers that block either the intracellular Ca^2+^ channels or extracellular Ca^2+^ channels. Asterisks denote significant change as observed by t-test.

The [Ca^2+^]_cyt_ inducibility of these miRNAs was further verified with the help of a Ca^2+^ ionophore i.e. A23187 that is expected to reverse the expression pattern obtained with inhibitors. A23187, also known as calcimycin, is an ionophore that binds to Ca^2+^ ions and acts as their carrier by inserting itself in the plasma membrane thereby acting as a channel for the divalent. Quantitative real time PCR profiling under control and ionophore treatment confirmed nine miRNAs (miR1425-5p, miR156a, miR159b, miR167h-5p, miR1862a,d, miR1876, miR1878 and miR396c-3p) that responded to channel inhibitors as well as to the ionophore but in inverse fashion, thereby confirming their [Ca^2+^]_cyt_ inducibility (Fig. 1c).

In order to identify the source of [Ca^2+^]_cyt_ inducibility of these miRNAs, subsequent analysis was done by treating rice seedlings with only internal or extracellular channel inhibitors to decipher the independent involvement of internal and external calcium stores in this regulation (Fig. 1d). Interestingly, miR156a, miR159b, miR167h-5p and miR396c-3p were significantly down-regulated when internal channels were inhibited but showed no remarkable change with the blocking of plasma membrane channels. On the other hand, miR166g-3p was down-regulated in the presence of external inhibitors only. Thus, specific miRNAs are uniquely regulated by the internal or external calcium reserves indicating toward the complexity of the regulatory phenomenon.

However from this data it appears that the effect of [Ca^2+^]_cyt_ is not global since only a small proportion of miRNAs was found to be responsive, indicative of transcriptional route (rather than miRNA processing where a higher proportion of miRNAs would have been differentially changed). Thus dissecting closely, the [Ca^2+^]_cyt_ responsiveness of the three main miRNA processing enzymes, namely DCL1, HYL1 and SE was checked and the response was found neither significant nor confirmed by ionophore treatment (Fig. S2).

Besides, we also found various calcium responsive *cis*-elements in the promoter regions of miRNAs. The promoter sequences of some miRNAs were subjected to a search for the calcium responsive motifs enlisted in literature (Galon et al., 2010). These motifs are ABA-related ABRE elements [(T/C)<u>ACGTG</u>(T/G)], CAMTA binding sites [(C/A)<u>CGCG</u>(T/G/C) and (C/A)<u>CGTGT</u>], [Ca^2+^]_cyt_ responsive elements [CARE-(C/A)<u>ACGTG</u>(T/G/C) and (C/A)<u>ACGCG</u>(T/G/C)], E-box [C<u>ANNTG</u>], G-box [C<u>ACGTG</u>], Z-box [AT<u>ACGTG</u>T], GT-box [GGTAATT], Rapid Stress Responsive Elements [<u>CGCG</u>TT]. For instance, miR1425 has 2 ABRE, 3 CAMTA and 5 E-box elements; miR156a has 1 CARE and 2 CAMTA sites and miR159b has 9 E-box sites.

### Targets of [Ca^2+^]_cyt_ regulated miRNAs in N22 are functionally widespread

To analyse the impact of these [Ca^2+^]_cyt_ regulated miRNAs, data from 8 degradome libraries including both in-house generated as well as publically available was analyzed to identify miRNA targets (Mutum *et al*., 2016). Targets identified under category ‘0-2’and ≥10 degradome reads were considered for the analysis (Table 1).

**Table. 1.** Targets of [Ca^2+^]_cyt_ responsive miRNAs in N22 as per degradome data. Targets having read no ≥10 and cat between 0-2 were considered.

| miRNA | Target loci | Target description | Read no | Category | P-value |
| --- | --- | --- | --- | --- | --- |
| miR1425-5p | LOC_Os10g35240 | Rf1, mitochondrial precursor | 280 | 0 | 0.0050 |
|  | LOC_Os10g35436 |  | 26 | 0 | 0.0054 |
|  | LOC_Os03g09110 | Mitochondrial Carrier protein | 11 | 0 | 0.0845 |
| miR156a | LOC_Os01g69830 | OsSPL2 - SBP-box gene family member | 67 | 0 | 0.0608 |
|  | LOC_Os02g04680 | OsSPL3 - SBP-box gene family member | 48 | 0 | 0.0132 |
|  | LOC_Os11g30370 | OsSPL19 - SBP-box gene family member | 45 | 0 | 0.0608 |
| miR159b | LOC_Os01g59660 | MYB family transcription factor | 172 | 0 | 0.0191 |
| miR160f-5p | LOC_Os06g47150, LOC_Os10g33940 | auxin response factor 18 | 1067 | 0 | 0.0262 |
|  |  |  | 1110 | 0 | 0.0262 |
|  | LOC_Os04g43910, LOC_Os02g41800 | auxin response factor | 51 | 0 | 0.0262 |
|  |  |  | 33 | 0 | 0.0262 |
| miR166g-3p | LOC_Os03g01890, LOC_Os10g33960 | START domain containing protein | 500 | 0 | 0.0242 |
|  |  |  | 123 | 0 | 0.0242 |
| miR167h-5p | LOC_Os02g06910 | auxin response factor 6 | 239 | 0 | 0.0277 |
|  | LOC_Os12g41950, LOC_Os06g46410 | auxin response factor | 46 | 0 | 0.0312 |
|  |  |  | 21 | 0 | 0.0312 |
|  | LOC_Os07g33790 | glutamate receptor 3.4 precursor | 27 | 2 | 0.4066 |

These targets included some well-known genes such as such as certain transcription factors like *OsSPL* (miR156a), *MYB* (miR159b and START domain containing (miR166), *auxin response factors* (miR160, miR167). A gene ontology classification of these targets (based on Molecular Function) shows transcription factors being the major category along with protein binding proteins and catalytic activity. The targets are widespread along almost all major categories like lipid and RNA-binding, kinase activity, transporters, structural molecule activity etc. (Fig. 2).

**Fig. 2.**
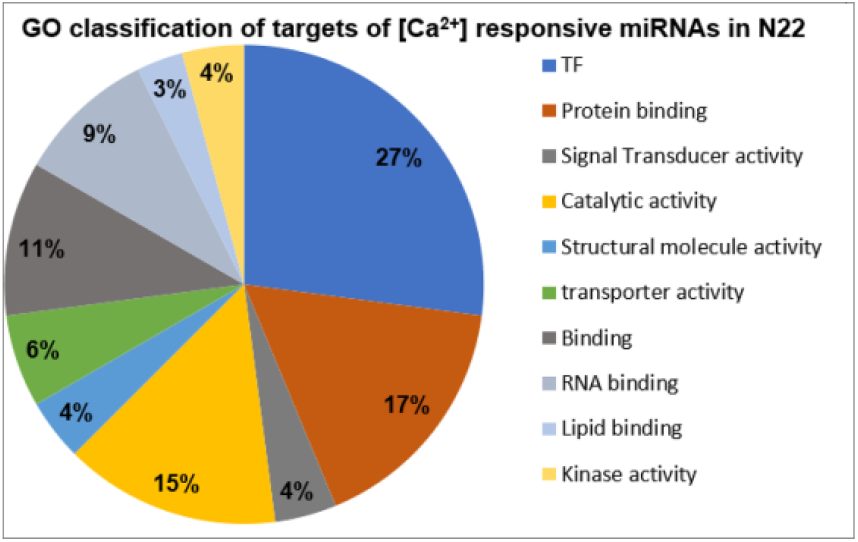
Classification of miRNA targets. Pie-chart representing different categories of miRNA-targets classified according to their GO term of ‘Molecular Function’.

Interestingly, upon relaxing the target identification criteria (cat:0-2 and read number ≥5) several new targets could be seen when searched against the Ricecyc database (Gramene) for possible involvement in metabolic pathways. Herein, miR1878 that is [Ca^2+^]_cyt_ responsive, seems to target Rubisco small subunit which is essential for Calvin cycle. Another interesting target enzyme that appears is NADH dehydrogenase (ubiquinone) involved in aerobic respiration (miR167h-5p) and a dehydrogenase involved in Reductive TCA cycle (miR156a) (Table.2).

**Table. 2.** [Ca^2+^]_cyt_ responsive miRNAs targeting genes involved in rice metabolic pathways. The targets are selected on basis of their cat: 0-2 along with read number ≥5

| miR | Target | Enzyme name | Pathway name | Read no | Category | P-value |
| --- | --- | --- | --- | --- | --- | --- |
| miR156a | LOC_Os02g38200 | Dehydrogenase | REDUCTIVE TCA CYCLE I | 8 | 2 | 0.632 |
|  | LOC_Os08g02700 | Fructosebiphosphate aldolase | XYLOSEMONOPHOSPHATE CYCLE | 9 | 2 | 0.788 |
|  | LOC_Os12g36194 | Nucleosidediphosphate kinase | DE NOVO BIOSYNTHESIS OF PYRIMIDINE DEOXYRIBONUCLEOTIDES | 5 | 2 | 0.918 |
| miR166g-3p | LOC_Os10g33800 | Malate dehydrogenase | MIXED ACID FERMENTATION | 13 | 2 | 0.973 |
| miR167h-5p | LOC_Os02g54254 | Lysineketoglutarate reductase | LYSINE DEGRADATION II | 5 | 2 | 0.996 |
|  | LOC_Os06g03830 | Oxidoreductase | ENTEROBACTIN BIOSYNTHESIS | 20 | 2 | 0.898 |
|  | LOC_Os07g45090 | NADH dehydrogenase (ubiquinone) | AEROBIC RESPIRATION ELECTRON DONOR III | 8 | 2 | 0.999 |
| miR396a-5p | LOC_Os01g01302 | Shikimate kinase | CHORISMATE BIOSYNTHESIS | 6 | 2 | 0.999 |
| miR1878 | LOC_Os12g19470 | Ribulosebiphosphate carboxylase | CALVIN CYCLE | 32 | 2 | 0.195 |

### Dehydration and ABA response of miRNAs is mediated by [Ca^2+^]_cyt_

Since it is a well-known fact that miRNAs respond to various abiotic stresses and that [Ca^2+^]_cyt_ plays a significant role in relaying these abiotic stress cues, we investigated whether the dehydration response of miRNAs in rice is mediated by [Ca^2+^]_cyt_. Thus, seven-day-old rice seedlings were subjected to air dehydration in presence or absence of calcium channel inhibitor cocktail as described previously. Subsequently, sRNA libraries were generated and sequenced (Supplementary Table. S2, S4). Comparison of both the datasets gives us an insight into [Ca^2+^]_cyt_ mediated dehydration response of miRNAs. Consequently, through qRT-PCR confirmation it was possible to identify that dehydration response of miR156a, miR167h-5p, miR168a-5p, miR5083 and miR5788 is being mediated by [Ca^2+^]_cyt_ levels (Fig 3a-c). Interestingly, miR156a and miR167h-5p also showed [Ca^2+^]_cyt_ responsiveness under control conditions.

**Fig. 3.**
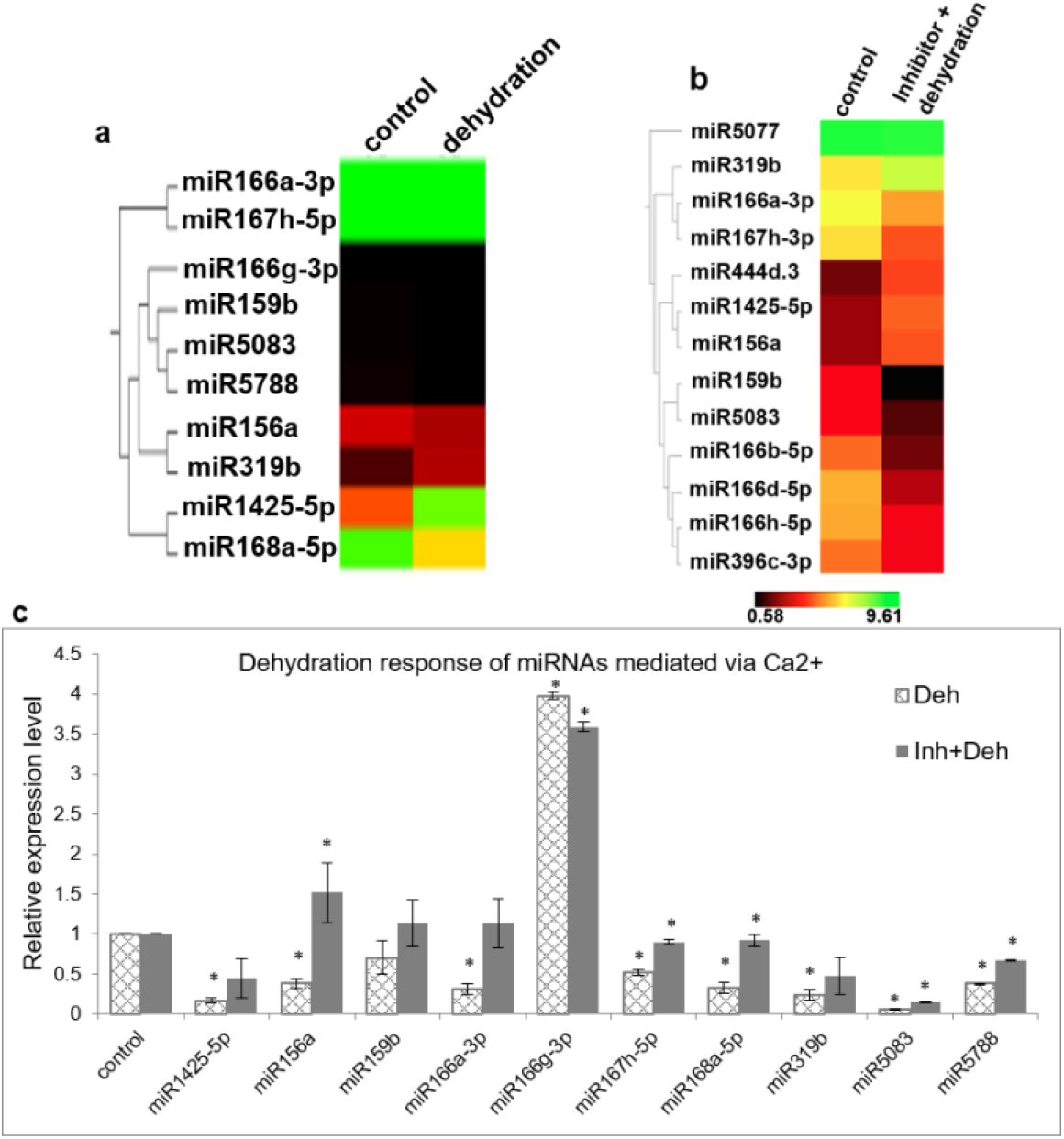
[Ca^2+^]_cyt_ mediates dehydration response of miRNAs in N22. Hierarchical clustering of miRNAs with significant fold change between **a** control and dehydrated seedlings; **b** control and seedlings treated with [Ca^2+^]_cyt_ channel inhibitors followed by dehydration; **c** Confirmation of calcium mediated dehydration response by qRT-PCR expression profiling of selected miRNAs. The asterisks denote significant change as observed by t-test.

Another important signalling molecule involved in regulation of gene expression is abscisic acid (ABA) aka the stress hormone that mediates several developmental processes in plants including dormancy, abscission, seed germination as well as abiotic and biotic stress response. ABA levels increase during certain stress conditions and mediate adaptive response. Thus, experiments were conducted to assess the response of miRNAs to elevated ABA levels by treating seven-days-old seedlings with ABA (100µM). Expression profiling of a selected set of miRNAs, based on previous knowledge, indicated that miR1425-5p, miR159b, miR168a-5p and miR529b are significantly down regulated while miR319b and miR530-5p are upregulated in presence of ABA (Fig 4).

**Fig 4.**
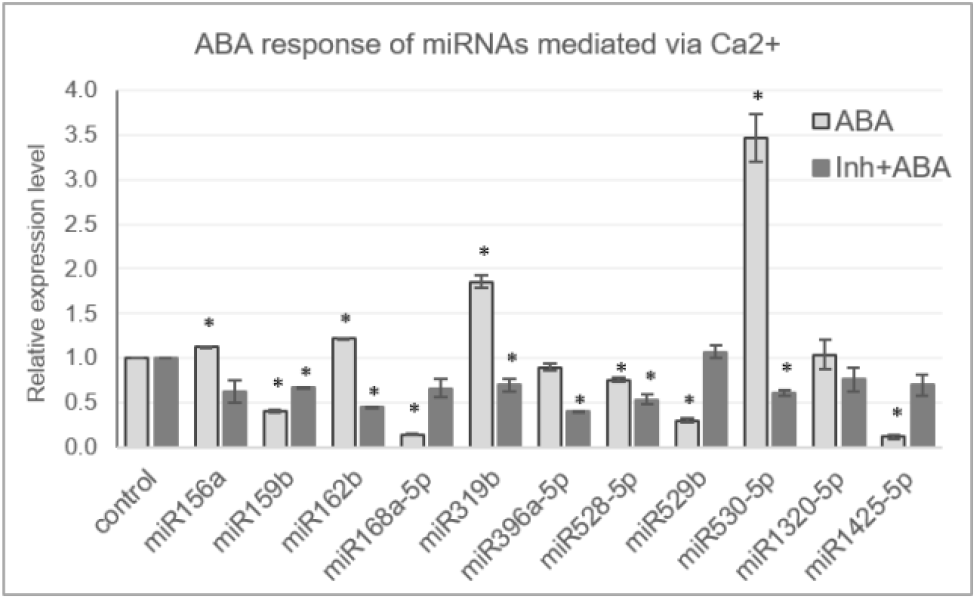
[Ca^2+^]_cyt_ mediates ABA response of miRNAs in N22. qRT expression profile of miRNAs under control, ABA (100 μM) and ABA pretreated with calcium channel inhibitors has been shown. The asterisks denote significant change as observed by t-test.

Expression of miR156a, miR162b, miR1320-5p, miR396a, miR528-5p did not deviate much from control. Correlation of the ABA and dehydration responsiveness indicate that miR1425-5p and miR168a-5p are downregulated during dehydration as well as ABA. miR156a is downregulated by dehydration but is not ABA responsive. On the other hand, miR162b, miR396a-5p, miR528-5p and miR1320-5p turn out neither dehydration nor ABA responsive. [Ca^2+^]_cyt_ was found to mediate ABA response of miRNAs as well, since pre-blocking [Ca^2+^]_cyt_ channels alleviated the hormone induced response of some miRNAs such as miR159b, miR319b, and miR530-5p (Fig 4). Notably, miR159b is responsive to [Ca^2+^]_cyt_ in resting state as well however its response to dehydration was not significant. Hence the demonstration of the versatile roles that [Ca^2+^]_cyt_ plays in regulating miRNAs under variable environmental conditions.

### Calmodulin and Calmodulin-binding Transcriptional Activators (CAMTAs) mediates the expression of [Ca^2+^]_cyt_ responsive miRNAs

[Ca^2+^]_cyt_ cues are perceived by sensor-relay proteins such as calmodulins that upon binding to [Ca^2+^]_cyt_ ions change their conformation and relay the signal by binding to other proteins. Thus as the next step, expression of [Ca^2+^]_cyt_ responsive miRNAs were checked for their dependency on calmodulin. To achieve this, rice seedlings were treated with 200 μM calmodulin inhibitor Trifluoperazine (TFP). As per qRT-PCR profiles, three miRNAs namely, miR156a, miR1878 and miR396c-3p are down regulated whereas miR1876, miR166g-3p, miR167h-5p and miR1425-5p were up-regulated upon calmodulin inhibitor treatment. Thus, CaM seems to positively relay the [Ca^2+^]_cyt_ signal to miR156a, miR1878, miR396c-3p, miR166g-3p and miR1425-5p while it appears to negatively affect miR1876 and miR167h-5p (Fig. 5).

**Fig. 5.**
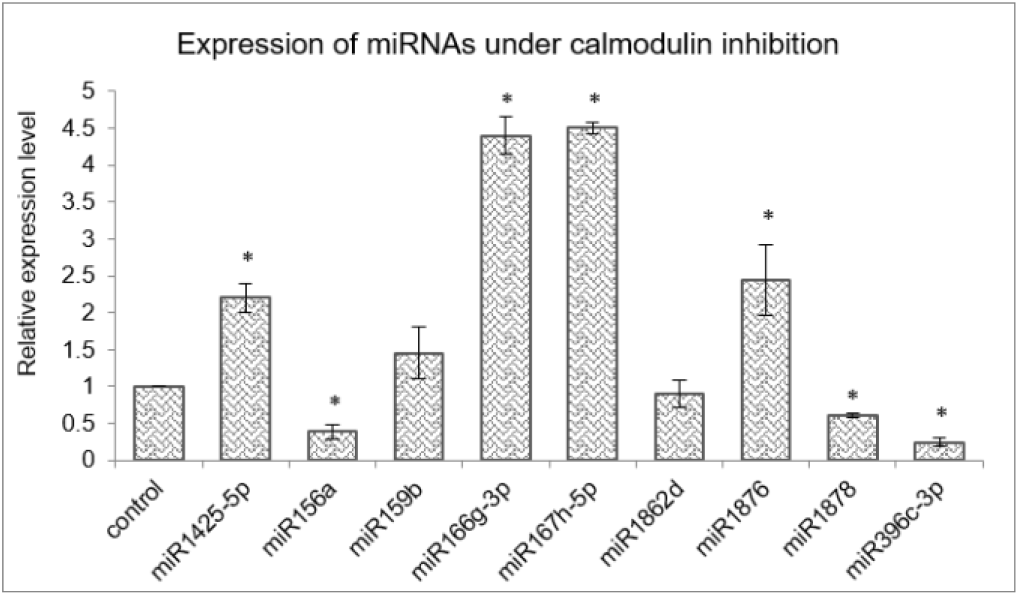
Calmodulins mediate miRNA expression in rice. Expression profiling to implicate calmodulins in mediating [Ca^2+^]_cyt_ response of miRNA genes in rice. The calmodulin inhibitor used is TFP (200 μM). The asterisk denotes significant change as observed by t-test.

Subsequent to finding [Ca^2+^]_cyt_ and calmodulin responsive miRNAs in rice, further components involved in mediating the [Ca^2+^] signal transduction of these miRNAs were explored. ‘Calmodulin-binding Transcriptional Activators’ or CAMTAs are calcium binding transcription factors that play an important role in mediating Ca^2+^/CaM mediated gene expression. Five miRNAs viz. miR156a, miR160a-5p, miR166a-3p, miR167h-5p and miR168a-5p are orthologous in both rice and *Arabidopsis* plus their promoter regions were found to have CAMTA binding sites. To investigate whether CAMTAs regulate their expression, these miRNAs were profiled in the *Arabidopsis camta* mutants (available at ABRC seed stock centre). There are six CAMTA genes in *Arabidopsis* and mutants specific to CAMTA loci *camta1, camta3, camta4, camta5* and *camta6* were procured (Supplementary Fig. 2).

Quantitative real time profiling for these miRNAs in the mutants revealed that miR156a is significantly reduced in *camta4* and *camta6* (Fig. 6). Expression of miR160a-5p is reduced in *camta4* while enhanced in *camta5* and *camta6*. Similarly, miR168a-5p is reduced in *camta3,4,5* but not in *camta6*. On the other hand, miR167h-5p is reduced in *camta5&6* but over accumulated in *camta1* mutant. Thus, these results indicate that CAMTAs are indeed involved in the regulation of miRNA expression. Promoter of miR167h has one CAMTA-binding site while that of miR156a harbours two sites. Further, CAMTA4 appears to have a broader and major role, as it appears to regulate the expression of all the five miRNAs. In case of miR156a, miR160a-5p, miR167h-5p and miR168a-5p it has a positive influence while for miR166a-3p it acts to negatively regulate it. Taking this cue further, we ventured to find if the orthologs of these AtCAMTA4 & 6 i.e OsCAMTA4 (LOC_Os04g31900) and OsCAMTA6 (LOC_Os07g43030) are involved in transcriptional regulation of miR156a and miR167h.

**Fig 6.**
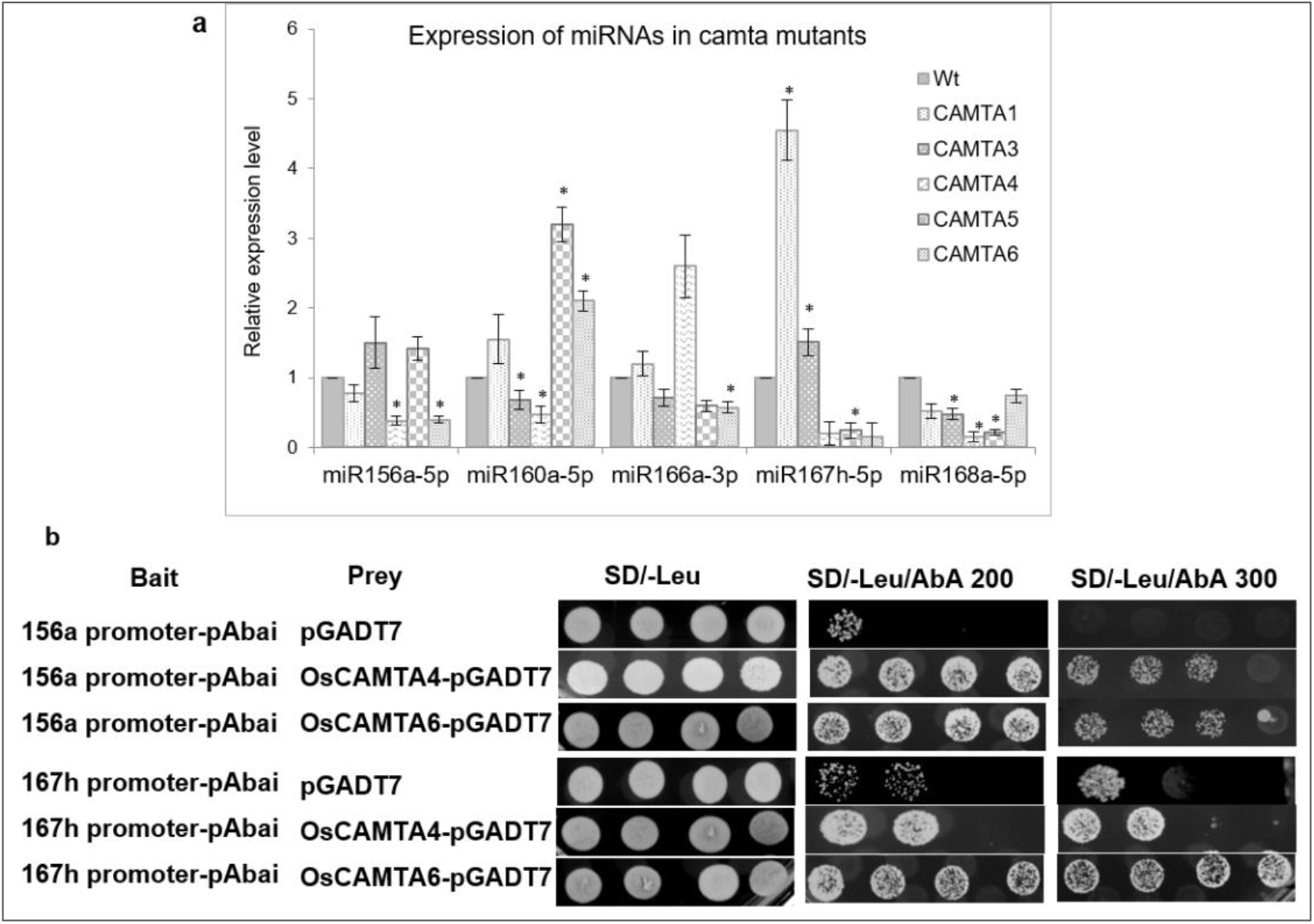
CAMTAs affect the expression of miRNAs. **a** qRT-PCR profile of miRNAs in wild type and the five *camta* mutant *Arabidopsis* lines. **b** Yeast-one-hybrid assay depicting the binding and trans-activation capability of OsCAMTA4 and OsCAMTA6 onto the promoters of miR156a and miR167h.

In order to find out whether OsCAMTA4 binds to the promoter of miR156a and miR167h, Y1H was performed using their promoter regions as bait. Promoter fragment containing the CAMTA-binding sites were cloned into the pABAi and transformed into Y1HGold to generate reporter strains. The basal expression of the bait reporter strain in the absence of prey was checked and found to be null. On the other hand, the full-length coding sequence of OsCAMTA4 & 6 were cloned into pGAD7 to generate prey vector. Both the prey vector as well as empty pGAD7 (as negative control) were transformed into Y1H gold strain harbouring promoter segments of miR156a and miR167h in separate experiments. Diploid yeast colonies containing miR156a promoter as bait and OsCAMTA4 & 6 as prey were able to grow strongly on the selective auxotrophic media containing aureobasidin A. But, miR167h shows a strong interaction with OsCAMTA6 and not with OsCAMTA4 (Fig6b). Thus hereby the physical interaction of the CAMTA-binding sites residing in the promoters of miRNA156a, miR167h with OsCAMTAs was confirmed in rice.

Furthermore, expression patterns of OsCAMTA4 & 6 were checked in different tissues under control and stress conditions. Both the CAMTAs respond to inhibitor treatment in seedlings by reduction in expression however their dehydration response could not be confirmed (Fig7a). In the mature drought stressed rice plant, both the miRNAs, miR156a and miR167h-5p are down regulated greatly in Flag Leaf while remaining close to control in spikelet (S. Balyan et al., 2017). Under the same drought conditions both the CAMTAs are downregulated significantly in the Flag Leaf with only OsCAMTA4 reducing in inflorescence (Fig 7c). Notably, these two miRNAs (miR156a, miR167h-5p) appear to co-regulate with OsCAMTA4 during seedling dehydration stress as well as FL drought stress. Thus, OsCAMTA4 appears to be a major regulator of these miRNAs under different growth and environmental stimuli.

**Fig 7.**
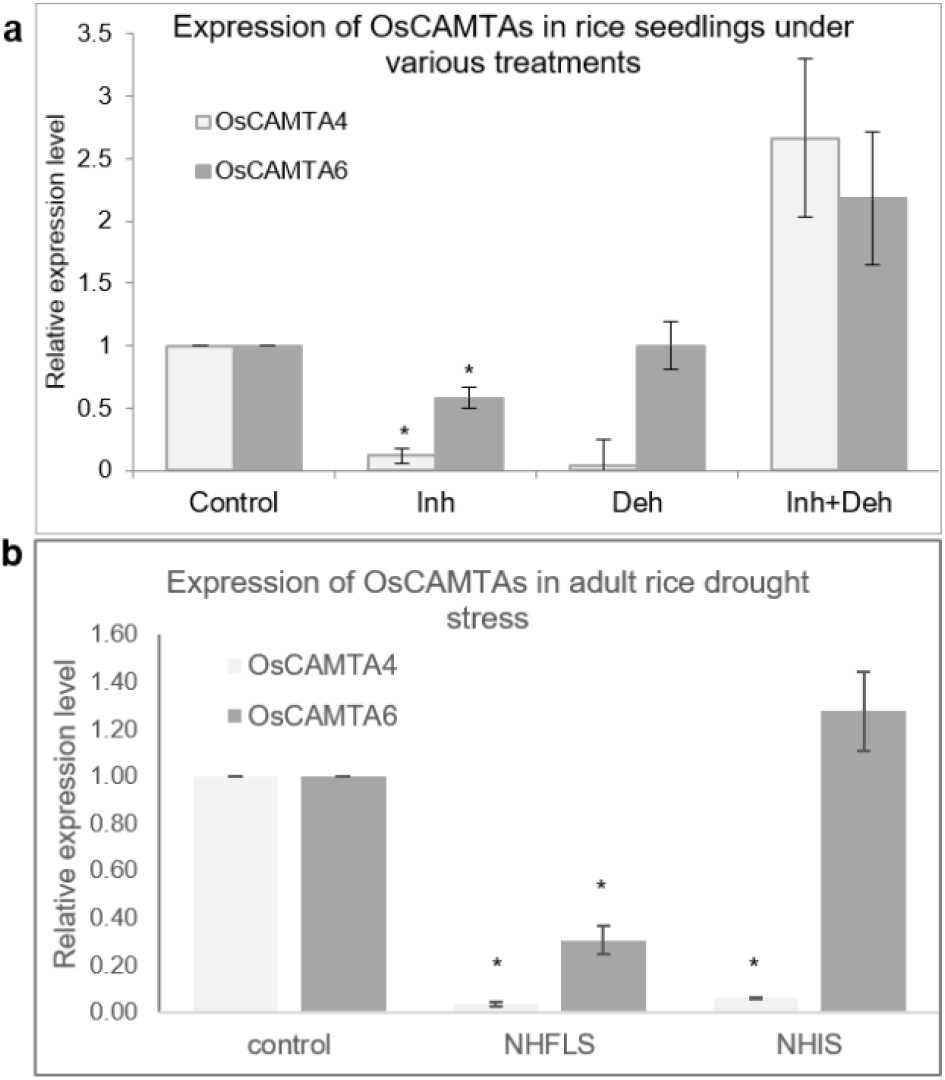
Expression profile of OsCAMTAs at different stages in rice. **a** Expression profile of *OsCAMTA4* and *OsCAMTA6* under [Ca^2+^]_cyt_ channel inhibition, dehydration and [Ca^2+^]_cyt_ channel inhibition followed by dehydration in N22 rice seedlings. **b** Expression profile of *OsCAMTA4* and *OsCAMTA6* under drought stress at heading stage as measured in the Flag Leaf and Inflorescence of N22 rice.

## DISCUSSION

These regulatory small molecules named as miRNAs are involved in almost all the critical biological processes in plants including growth and development as well as combating various stresses. While significant number of studies have been done to understand the molecular and biochemical processes that are regulated by miRNAs, few studies attempt to study the regulation of miRNA genes themselves. To address this area, we investigated the molecular nature of the schema that regulates miRNA genes by looking into the involvement of one of the most important signaling molecule i.e. cytosolic levels of calcium in regulating miRNA gene expression. In the past, Kaplan et al., 2006 studied the effect of [Ca^2+^]_cyt_ bursts induced by calmodulin inhibitors on gene expression of protein coding genes in *Arabidopsis*. Over 200 genes were found to be differentially regulated (162 up regulated and 68 down regulated significantly) and it was summarized that [Ca^2+^]_cyt_ bursts in the cytosol affect the transcriptome to a certain extent and is responsible for induction of signaling related genes. In another study where different types of [Ca^2+^]_cyt_ bursts were given (single peak, oscillations and prolonged [Ca^2+^]_cyt_ elevation) by electrical stimulus, changes in gene expression were observed in *Arabidopsis* (Whalley et al., 2011). Herein again, the authors found more number of genes up regulated than down regulated in response to [Ca^2+^]_cyt_ single peak burst (104 up vs 30 down regulated genes) and oscillations (256 up regulated vs 97 down regulated genes) with a large number of overlapping genes among the up-regulated ones. In our study as well, we found that miRNAs indeed responding to [Ca^2+^]_cyt_ levels. The proportion of such miRNAs in N22 (∼2.8%) is pretty close to the estimate of protein coding genes found by Kaplan *et al* in *Arabidopsis* (∼3%) and by Feske et al., 2001 in human T lymphocyte cells (∼2.1%). Additionally, higher proportion of miRNAs was up regulated by [Ca^2+^]_cyt_ than down regulated which is similar to the findings as mentioned above. Among the [Ca^2+^]_cyt_ responsive miRNAs as well, a big proportion of miRNAs (such as miR156, miR159, miR166, miR167, miR319, miR396) are also responsive to at least one or multiple abiotic stresses such as cold, drought, salt, UV and mechanical (Balyan et al., 2015).

The data described above identified several miRNAs whose expression is regulated by cytosolic calcium levels. On the other hand, it is also possible that miRNAs target genes involved in the calcium signaling cascade. Although prediction based models have specified [Ca^2+^]_cyt_ signaling components as miRNA targets for a long time now (Sunkar et al., 2008), our study identifies those targets on the basis of degradome data. Data analysis identified genes such as IQ CaM-binding motif family protein (miR531b), OsCML16 (miR5517), OsCML22 (miR164a,c,d,e), CAMK_CAMK_like.9 (miR2090), CAMK_CAMK_like.11 (miR5151), MAPKKK (miR1428e-5p, miR2876-3p). In an earlier study in tomato, an autoinhibited [Ca^2+^]_cyt_ ATPase *ACA10* has been shown to be targeted by miR4376. This miR-target module is involved in proper fruit development since the overexpression of either the miRNA or the miR-resistant ACA10 resulted in elongated stamen filaments as well as reduced number of mature fruits (Wang et al., 2011). Thus, it appears that miRNAs play an important role in calcium mediated regulatory schema of the plant cell.

[Ca^2+^]_cyt_ is known to be the mediator of several biotic and abiotic stress responses like heat, drought, touch, cold and salinity (Albrecht et al., 2003; Choi et al., 2005; Knight et al., 1996; Schulz et al., 2013; B. Wu et al., 2012). In this study, we show that [Ca^2+^]_cyt_ also mediates the dehydration response of several miRNA genes as well. These included miRNAs like miR156a, miR167h-5p, miR168a-5p and miR5788. Among these, the former two are also responsive to resting [Ca^2+^]_cyt_ under non-stress conditions while latter two are not. The trend exhibited by the latter group could be speculated by the involvement of some other factors such as dehydration specific transcription factors that might be controlled by [Ca^2+^]_cyt_ signaling. Earlier, H_2_S which is a potent signaling messenger in various plant physiological processes including drought (Jin et al., 2011, 2013), has been shown to mediate drought response of some miRNAs like miR167a/c/d, miR393a, miR396a, miR398a/b/c in *Arabidopsis* (Jiejie Shen et al., 2013). Another significant hormone known to be involved in many abiotic stresses especially drought/dehydration stress-the abscisic acid and its influence on miRNA expression-has also been discussed in this study. In general, [Ca^2+^]_cyt_ is known to mediate many ABA responses such a stomatal closure and seedling growth (Y. Guo et al., 2002; McAinsh et al., 1990) and thus there is a significant overlap of ABA and calcium mediated signalling. Indeed, expression of several miRNAs was found to be regulated by ABA. Expression trends of miR1425-5p, miR156a, miR162b, miR168a, miR319b, miR528-5p, miR530-5p matches with previous known knowledge while rest had no previous information regarding their ABA response (Shen et al., 2010; Tian et al., 2015). Further analysis conducted to decipher whether modulation in the cytosolic calcium levels mediate ABA inducibility of the miRNAs resulted in the finding that cytosolic calcium levels do indeed mediate the ABA response of miR159b, miR319b, miR530-5p.

Further investigation into the signaling cascade regulating the miRNA expression revealed the involvement of calmodulins (miR156a, miR1878, miR396c-3p, miR166g-3p, miR167h-5p, miR1425-5p, miR1876). Calmodulins are signal sensor-relay proteins that directly bind to calcium and relay the signal to other proteins such as CAMTAs (calmodulin binding transcription activators) that bind to particular motifs in promoter sequences of genes and either activate or suppress transcription. The use of calmodulin inhibitors and *camta* knock-down mutants affected the expression of certain miRNAs namely, miR156a, miR160a-5p, miR166a-3p, miR167h-5p and miR168a-5p. Besides, several calcium responsive promoter motifs have been identified in the promoter sequences of calcium responsive genes (Finkler et al., 2006; Galon et al., 2010; Kaplan et al., 2006). Accordingly, we found the presence of these calcium responsive motifs namely, CARE (calcium responsive elements), ABRE (ABA responsive elements and related motifs), E-box, G-box, GT-box, Z-box and CAMTA binding sites in the promoter sequences of calcium responsive miRNAs. The Y1H shows strong physical interaction between OsCAMTA4 and miR156a but this CAMTA does not interact with miR167h. However, OsCAMTA6 shows strong interaction with the promoters of miR156a and miR167h. Thus, hereby we reveal novel and key players in the transcriptional regulation of miR156a and miR167h. While miR156a is seen to be responsive to [Ca^2+^]_cyt_ in resting conditions, positively by calmodulin as well as [Ca^2+^]-mediated dehydration, miR167h-5p has a slightly different story. Although [Ca^2+^]_cyt_ affects its expression under control conditions as well as dehydration, miR167h-5p shows negative regulation by calmodulins. Both of these miRNAs show binding and trans-activation by OsCAMTA4 who appears to be co-regulated in seedling dehydration as well as FL drought stress. This appears as a significant finding since miRNA expression has till date not been associated with CAMTAs.

miR156 has been studied in quite details and there is plenty of knowledge available about its function and regulation. The miRNA is a floral repressor and promoter of juvenile phase in *Arabidopsis* (Wu et al., 2009) is involved in lateral root development and leaf morphology in Arabidopsis (Gao et al., 2018). Regarding its regulation, it has been shown that AGL15 and AGL18 act in cooperation to promote its transcription by binding to the CArG motifs in its promoter (Serivichyaswat et al., 2015). The Phytochrome –Interacting Factors or PIF1, PIF3, PIF4 and PIF5 have also been shown to directly bind and repress the expression of miR156b/d/e/f/h to enhance the shade avoidance syndrome in *Arabidopsis*. Another protein called DOG1 (DELAY OF GERMINATION 1) plays a role in efficient processing of primary miR156 to its active mature form by regulating the processing proteins-DCL1, HYL1, SE, TGH and CDC5 (Huo et al., 2016). Recently, another player was found in miR159 that targets MYB33 that in turn binds and promotes the transcription of miR156a&c during the young seedling stage (C. Guo et al., 2017). Furthermore, clues behind its temporal expression pattern have been found in epigenetic regulation. The transcription activating mark H3K4me3 is seen abundantly at the miR156a and miR156c loci during the early seedling stages wherein it contributes to its high expression (Y. Xu et al., 2018) while during vegetative phase change there is an increase in the level of histone H3K27me3 with simultaneous decrease in H3K4me4 and H3K27ac at regions upstream as well as immediately downstream of its TSS resulting in its decline in abundance (Xu et al., 2016). Besides, a cycling DOF transcription factor CDF2 has also been shown to be its transcriptional activator (and repressor of miR172) acting in the same signaling pathway to control flowering in *Arabidopsis* (Sun et al., 2015). Regarding the regulation of miR156 in rice, drought has already been shown to down regulate the miRNA in inflorescence tissue (L. Zhou et al., 2010). Besides, it is also responsive to several hormones such as auxin in *Arabidopsis* [down regulation; (Marin et al., 2010)] and ethylene in tomato [down regulation; (Zuo et al., 2012)]. In rice, the miRNA has been shown to be independent of any regulation by gibberellin during juvenile to adult phase transition (Tanaka, 2012). Our data brings more regulators of miR156 into light which are [Ca^2+^] and OsCAMTA4 & 6. Since these are located pretty high in the signalling hierarchy that generates any and all responses inside a cell, this new mode of regulation might help to explain the various responses miR156 displays under the various abiotic stresses and different stages of the lifecycle of the plant.

miR167 is also a conserved miRNA across monocots and dicots and is known to target the ARF6/8 genes thereby acting as a node in auxin signaling (Barik et al., 2015). It is known to be involved in adventitious root development in rice (Meng et al., 2011), modulation of auxin signaling during bacterial infection in tomato (Jodder et al., 2017), in regenerating calli in rice (Sinha et al., 2019), in blue-light signaling in *Arabidopsis* (Pashkovskiy et al., 2016) and abiotic stress such as salinity (Ding et al., 2009; Frazier et al., 2011; Liu et al., 2008), drought in rice (S. Balyan et al., 2017). For such an evolutionary conserved and functionally significant miRNA, the information about its own regulation was lacking. In our study we showed its regulation via [Ca^2+^]_cyt_, calmodulin and OsCAMTA6. Its dehydration response is also mediated via [Ca]^2+^_cyt_. Thus, it’s a remarkable discovery for a miRNA that acts as a node in a critical signaling pathway such as the auxin signaling ultimately regulating a plethora of plant functions.

## Conclusions

The understanding of the regulatory pathways governing the expression of another class of regulators i.e. the miRNAs is critical in order to be able to manipulate them for advantageous traits in plants. With this view the study explored the involvement of cytosolic calcium in regulating expression of miRNA genes under control and drought stress conditions that we could clearly establish with the help of calcium channel inhibitors and ionophore. The fact that [Ca]^2+^ mediates several abiotic responses was also explored and it was demonstrated it is the same for miRNAs in rice as well. The further dissection revealed the involvement of calmodulins and CAMTAs. Through yeast-one-hybrid experiments, OsCAMTA4 & 6 were proved to bind the CAMTA binding sites of the very critical miR156a and miR167h. Both these CAMTAs were found to be coregulated with these two miRNAs at various developmental and stress stages indicative of a possible regulatory schema for these miRNAs in rice.

## Declarations

## Funding/Acknowledgements

The study was funded by Science and Engineering Research Board, Department of Science and Technology under the Grant no: EMR/2016/006081 awarded to the corresponding author. SK is grateful to CSIR for fellowship and RDM and VP thank Delhi University for the fellowships during the research tenure.

## Author Contributions

SR conceived the concept and SR and SK designed the experiments. SK constructed the NGS libraries and performed the data analysis and qRT-PCR. VP performed qRT-PCR and yeast-one-hybrid experiments. RDM performed the degradome analysis. SK and SR prepared the manuscript. All authors read and approved the final manuscript.

## Availability of data and material

The datasets used and/or analysed during the current study are available from the corresponding author on reasonable request. All data generated or analysed during this study are included in this published article [and its supplementary information files].

## Compliance with Ethical Standards/Conflict of interest

Authors declare no conflict of interest.

## Ethics approval

NA

## Consent to participate

NA

## Consent for publication

All authors consent to publish.

## Notes

### Competing Interest Statement

The authors have declared no competing interest.

