## Supplementary Figures for "Investigations on regulation of miRNAs in rice reveal [Ca^2+^]_cyt_ signal transduction regulated miRNAs"

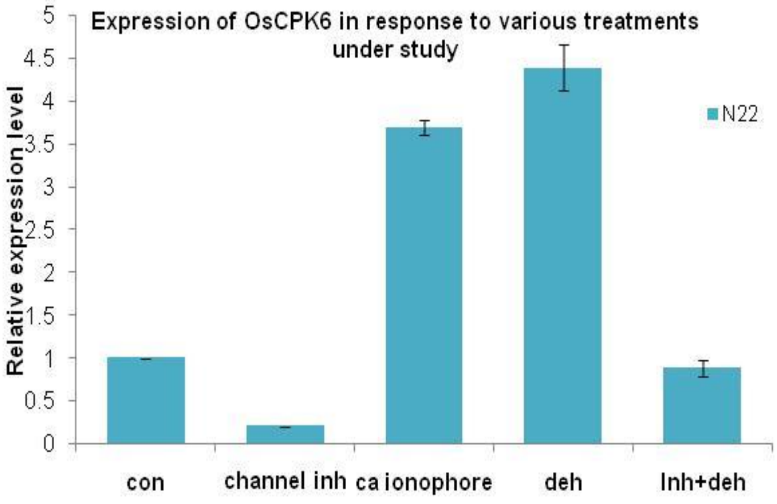


Fig S1: Expression of OsCPK6 in response to various treatments under study

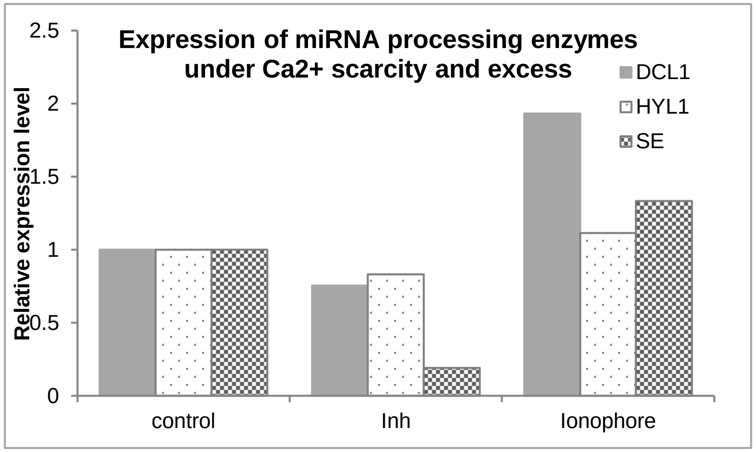


Fig S2: Expression of *DCL1, HYL1 and SE* under calcium channel inhibition and excess.

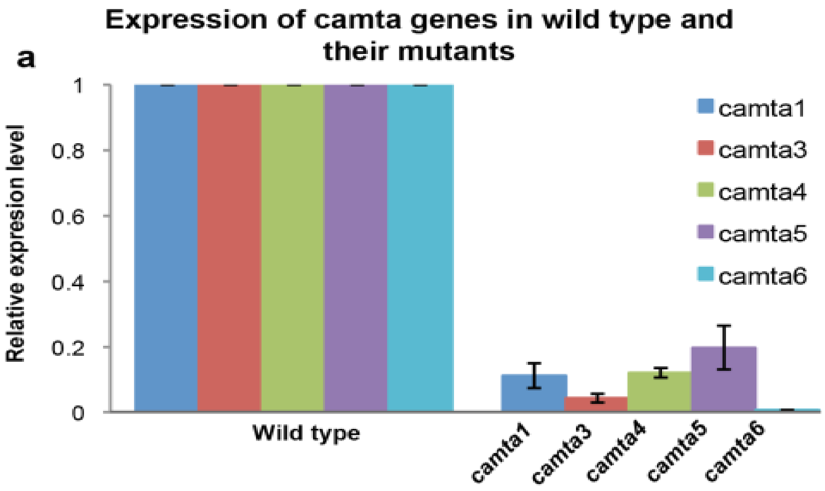


Fig S3: Expression of At*CAMTA* genes in their respective mutants
