## Supplementary Tables for "Investigations on regulation of miRNAs in rice reveal [Ca^2+^]_cyt_ signal transduction regulated miRNAs"

**Supplementary Table S1: List of primers used in the study**

| Primer ID | Sequence |
|---|---|
| At5S-RT-Forward | CGATGAAGAACG TAGCGAAATG |
| os-5s-forward | TGCGATCATACCAGCACTAAAGC |
| osa-miR1320-5p | CGATGGAACGGAGGAATTTTATAG |
| osa-miR1425-5p | CTAGGATTCAATCCTTGCTGCT |
| osa-miR156a | CGTGACAGAAGAGAGTGAGCAC |
| osa-miR159b | TTTGGATTGAAGGGAGCTCTG |
| osa-miR159f | CCTTGGATTGAAGGGAGCTCTA |
| osa-miR160f-5p | TGCCTGGCTCCCTGAATGCCA |
| osa-miR162b | TCGATAAGCCTCTGCATCCAG |
| osa-miR166a-3p | TCGGACCAGGCTTCATTCCCC |
| osa-miR166g-3p | TCGGACCAGGCTTCATTCCTC |
| osa-miR166e-3p | TCGAACCAGGCTTCATTCCCC |
| osa-miR167h-5p | TGAAGCTGCCAGCATGATCTG |
| osa-miR168a-3p | GATCCCGCCTTGCACCAAGTGAAT |
| osa-miR168a-5p | GCTTGGTGCAGATCGGGAC |
| osa-miR1861j | AGGTCTTGAGGCAGGAACTGAG |
| osa-miR1862a-d | GACTAGGTTTGTTTATTTTGGGACG |
| osa-miR1876 | GGTTTGTGGGCTGGCCC |
| osa-miR1878 | GCGCACTTAATCTGGACACTATAAAAGA |
| osa-miR319b | TTGGACTGAAGGGTGCTCCC |
| osa-miR396a-5p | TTCCACAGCTTTCTTGAACTG |
| osa-miR396c-3p | AGGTCAAGAAAGCTGTGGGAAG |
| osa-miR444b.1 | TGTTGTCTCAAGCTTGCTGCC |
| osa-miR5072 | CGATTCCCCAGCGGAGTCGCCA |
| osa-miR5788 | TGGATGTGACATACTCTAGTA |
| osa-miR5794 | TGAGGAATCACTAGTAGTCGT |
| osa-miR5083 | GCGCGAGACTACAATTATCTGATCA |
| osa-miR528-5p | TGGAAGGGGCATGCAGAG |
| osa-miR529b | CGCAGAAGAGAGAGAGTACAGCTT |
| osa-miR530-5p | 5'TGCATTTGCACCTGCACCTA3' |
| probe Fam at 5' end and BHQ at 3 | 5'{6FAM}CTTGGTACCGAGCTCGGATCCACTAGTCC {BHQ1}3 |
| RTQ_UNI_REV_NEW | 5' CGAATTCTAGAGCTCGAGGCA3' |
| AtCAMTA1-RT-F | CTGTCAGAAGCCCAACACAG |
| AtCAMTA1-RT-R | CCTTGAGCTTCTCATGAGCTTCTC |
| AtCAMTA3-RT-F | CAACGACATCCAAGAAAGCA |
| AtCAMTA3-RT-R | TGAGGACATAGGCAACATCAA |
| AtCAMTA4-RT-F | TTTGGAAAGGGCAGGAACTA |
| AtCAMTA4-RT-R | TTTGGTAACCTCGCACATGA |
| AtCAMTA5-RT-F | ATCGCGAGACACATGAGGTT |
| AtCAMTA5-RT-R | GACTGTTGCTCCGCACT |
| AtCAMTA6-RT-F | TGGAGTCTACCGTTGCATCA |
| AtCAMTA6-RT-R | TTGTCTTCAGGGACGGTCTT |
| OsCAMTA4-RT-F | CCTCGTCGGACCACTTGGT |
| OsCAMTA4-RT-R | TCCTCCGTCGAGCCTGAA |
| OsCAMTA6-RT-F | CAAGTACGGGCTGCCCAAT |
| OsCAMTA6-RT-R | GGACGTAGCCATCGATTCTGA |
| OsCAMTA4-Y1H-F | CCCGGGATGAGCCTGAGTTTT |
| OsCAMTA4-Y1H-R | CTCGAGCTATTCAGCAGTGGC |
| OsCAMTA6-Y1H-F | CCCGGGATGGCGGAGGTTCGCAAGTACG |
| OsCAMTA6-Y1H-R | GAGCTCTCACAAAATGGTGGGCATTGGTGCG |

| | |
|---|---|
| m156a-promoter-Y1H-F | GAGCTCTGCATATTTTGTCAGCCT |
| m156a-promoter-Y1H-R | CTCGAGACAAATGATCCATGCAA |
| m168a-promoter-Y1H-F | CGAGCTCACCAGGCCTAAACAC |
| m168a-promoter-Y1H-R | CCGCTCGAGTACCACAACAAGACG |
| m167h-promoter-Y1H-F | AAGCTTCTGTAGTGCAAGCCTTGCACGCC |
| m167h-promoter-Y1H-R | CTCGAGCACCTGTCTGGTAGCTGGCGTATTGTG |

Table S2: Details of the small RNA libraries generated for the various treatments given to 1 week old N22 rice seedlings

| | Control | | Calcium channel | | Dehydration | | $Ca^{2+}$ channel | |
|---|---|---|---|---|---|---|---|---|
| **Total number of reads** | 38,42,993 | 47,52,109 | 44,77,081 | 62,32,823 | 51,15,041 | 55,99,592 | 56,82,314 | 17,40,671 |
| **Number of reads post trimming** | 22,18,013 | 22,21,756 | 20,15,332 | 15,75,311 | 35,87,579 | 42,29,078 | 30,49,540 | 4,33,949 |
| **Number of miRNAs detected** | **293** | **303** | **258** | **279** | **317** | **313** | **357** | **214** |

| Table S3. NGS data of miRNAs showing significant differential response to calcium channel inhibitors in N22 | | |
|---|---|---|
| **Small RNA - Name** | **test: control vs inhibitors, tagwise dispersions - P-value** | **test: control vs inhibitors, tagwise dispersions - Fold change** |
| osa-miR1425-5p | 0.0326 | 3.8445 |
| osa-miR156ab-5p/c-5p/d/e/f-5p/g-5p/h-5p/6i/j-5p | 0.0284 | -1.9567 |
| osa-miR159a.1//osa-miR159b | 0.0421 | -4.8172 |
| osa-miR160f-5p | 0.0489 | 25.9825 |
| osa-miR166g-3p//osa-miR166h-3p | 0.0272 | 6.3279 |
| osa-miR167d-5p/e-5p/f/g/h-5p/i-5p/j | 0.0165 | -31.6562 |
| osa-miR168a-3p | 0.0209 | 31.1884 |
| osa-miR1862a/b/c | 0.0176 | -4.3178 |
| osa-miR1862d | 0.0265 | -3.5770 |
| osa-miR1876 | 0.0029 | -4.9732 |
| osa-miR1878 | 0.0348 | -5.5072 |
| osa-miR396a-5p/b-5p/6c-5p | 0.0117 | 7.3611 |
| osa-miR396c-3p | 0.0361 | -4.6232 |
| osa-miR444b.1/c.1 | 0.0414 | 2.2750 |
| osa-miR5072 | 0.0017 | 3.6347 |
| osa-miR5788 | 0.0021 | -13.3537 |
| osa-miR5794 | 0.0170 | -2.2896 |
| osa-miR159f | 0.87532 | 1.124644 |
| osa-miR166e-3p | 1 | 1.069284 |
| osa-miR1320-5p | 0.1764 | -3.45089 |
| osa-miR319a-3p.2-3p//osa-miR319b | 0.468171 | 1.61154 |
| osa-miR529b | 1 | 1.105139 |

| Table S4a. NGS data of miRNAs showing significant differential response to dehydration in N22 | | |
|---|---|---|
| **Small RNA - Name** | **EDGE test: con vs deh - P-value** | **EDGE test: con vs deh- Fold change** |
| osa-miR1425-5p | 0.0110 | 1.4110 |
| osa-miR156a/b-5p/c-5p/d/e/f-5p/g-5p/h-5p/i/j-5p | 0.0241 | -1.7050 |
| osa-miR159a.1/b | 0.0062 | -4.9017 |
| osa-miR164e | 0.0203 | 28.6170 |
| osa-miR166a-3p/b-3p/c-3p/d-3p/f/m/j-3p | 0.0059 | -2.0315 |
| osa-miR166g-3p/h-3p | 0.0174 | -5.5461 |
| osa-miR166j-5p | 0.0036 | 4.3963 |
| osa-miR166k-3p/166l-3p | 0.0114 | -1.4979 |
| osa-miR167d-5p/e-5p/f/g/h-5p/i-5p/j | 0.0163 | -1.5456 |
| osa-miR167h-3p | 0.0049 | -2.8318 |
| osa-miR168a-5p | 0.0054 | -2.0747 |
| osa-miR169f.2 | 0.0017 | 5.6306 |
| osa-miR169i-5p.2 | 0.0494 | 2.1616 |
| osa-miR171b/c-3p/d-3p/e-3p/f-3p | 0.0016 | 2.9817 |
| osa-miR171e-5p | 0.0264 | -4.2242 |
| osa-miR1861c/e/k/m | 0.0304 | 7.0910 |
| osa-miR1862d | 0.0018 | 42.4347 |
| osa-miR1876 | 0.0340 | -1.8862 |
| osa-miR2873a | 0.0425 | -24.8667 |
| osa-miR319a-3p.2-3p/b | 0.0386 | 1.6711 |
| osa-miR394 | 0.0111 | 32.1169 |
| osa-miR396a-5p/b-5p | 0.0400 | 2.7115 |
| osa-miR396c-3p | 0.0327 | -2.0362 |
| osa-miR396c-5p | 0.0259 | 1.5672 |
| osa-miR396e-5p | 0.0041 | 3.2247 |
| osa-miR3979-3p | 0.0063 | 8.5728 |
| osa-miR5083 | 0.0437 | -3.0758 |
| osa-miR5144-5p | 0.0342 | 3.8935 |
| osa-miR5150-3p | 0.0427 | 2.3370 |
| osa-miR5150-5p | 0.0012 | 2.4879 |
| osa-miR529a | 0.0467 | 2.3693 |
| osa-miR535-5p | 0.0012 | 12.3303 |
| osa-miR5513 | 0.0063 | 3.9353 |
| osa-miR5788 | 0.0226 | -3.4520 |
| osa-miR6249a/b | 0.0236 | -3.9106 |
| osa-miR812o-3p | 0.0203 | 28.6170 |

**Table S4b. NGS data of miRNAs showing significant differential response to inhibitor+dehydration in N22**

| Small RNA - Name | EDGE test: con vs inh-deh- P-value | EDGE test: con vs inh-deh- Fold change |
|---|---|---|
| osa-miR1425-5p | 0.0103 | 4.4578 |
| osa-miR156a/b-5p/c-5p/d/e/f-5p/g-5p/h-5p/i/j-5p | 0.0421 | -1.1536 |
| osa-miR159a.1//osa-miR159b | 0.0231 | -2.1801 |
| osa-miR166a-3p/b-3p/c-3p/d-3p/f/m/j-3p | 0.0456 | -2.1259 |
| osa-miR166b-5p | 0.0242 | -5.5970 |
| osa-miR166d-5p | 0.0102 | -5.3681 |
| osa-miR166h-5p | 0.0209 | -3.5214 |
| osa-miR167h-3p | 0.0298 | -2.8009 |
| osa-miR319a-3p.2-3p/b | 0.0492 | 2.0217 |
| osa-miR396c-3p | 0.0449 | -2.7098 |
| osa-miR444d.3 | 0.0462 | 4.4051 |
| osa-miR5077 | 0.0192 | -2.3474 |
| osa-miR5083 | 0.0238 | -4.0065 |
